# A green tea polyphenol Epigallocatechin-3-gallate modulates Tau Post-translational modifications and cytoskeletal network

**DOI:** 10.1101/2020.03.22.002014

**Authors:** Shweta Kishor Sonawane, Subashchandrabose Chinnathambi

## Abstract

**Background:** Alzheimer’s disease is a type of dementia denoted by progressive neuronal death due to the accumulation of proteinaceous aggregates of Tau. Post-translational modifications like hyperphosphorylation, truncation, glycation, *etc*. play a pivotal role in Tau pathogenesis. Glycation of Tau aids in paired helical filament formation and abates its microtubule-binding function. The chemical modulators of Tau PTMs, such as kinase inhibitors and antibody-based therapeutics, have been developed, but natural compounds, as modulators of Tau PTMs are not much explored.

**Methods:** We applied biophysical and biophysical techniques like fluorescence kinetics, SDS-PAGE, western blot analysis and transmission electron microscopy to investigate the impact of EGCG on Tau glycation *in vitro*. The effect of glycation on cytoskeleton instability and its EGCG-mediated rescue were studied by immunofluorescence in neuroblastoma cells.

**Results:** EGCG inhibited methyl glyoxal (MG)-induced Tau glycation *in vitro*. EGCG potently inhibited MG-induced advanced glycation endproducts formation in neuroblastoma cells as well modulated the localization of AT100 phosphorylated Tau in the cells. In addition to preventing the glycation, EGCG enhanced actin-rich neuritic extensions and rescued actin and tubulin cytoskeleton severely disrupted by MG. EGCG maintained the integrity of the Microtubule Organizing Center (MTOC) stabilized microtubules by Microtubule-associated protein RP/EB family member 1 (EB1).

**Conclusions:** We report EGCG, a green tea polyphenol, as a modulator of *in vitro* methylglyoxal-induced Tau glycation and its impact on reducing advanced glycation end products in neuroblastoma cells. We unravel unprecedented function of EGCG in remodeling neuronal cytoskeletal integrity.

## Introduction

Tau is a microtubule-associated protein, which aids in neuronal functioning (1,2) and Tau neurofibrillary tangles is one of the important characteristic pathology in AD (3,4). Aggregation of Tau leads to loss of its physiological function and adds up to the pathology of AD (5,6). Tau undergoes various PTMs that either leads to its aggregation or prevent it. The pro-aggregation PTMs of Tau include phosphorylation (7-10), truncation (11,12), glycation (13), acetylation (14,15), sumoylation (16,17), ubiquitination (18,19) and nitration (20-22). The anti-aggregation PTMs include O-glycosylation (23,24), dephosphorylation (25,26) and prolyl isomerization (27,28). The cross talk between these pro and anti-aggregation PTMs has an impact on Tau pathology (29). The clearance of hyperphosphorylated Tau is decreased by PTMs like glycation, nitration and polyamination whereas, glycosylation and dephosphorylation prevents Tau hyperphosphorylation.

Glycation, unlike phosphorylation is a non-enzymatic PTM, occurring between reducing sugars and protein, lipids etc. Glycation is triggered in presence of high blood sugar levels due to their metabolism *via* polyol pathway, which converts sugars into highly reactive intermediates like methyl glyoxal (MG), gloxal *etc*. Glycation involves multistep reactions including complex re-arrangements forming advanced glycation end products (AGEs). AGEs cause protein cross-linking hampering its function (30). AGEs play a pathological role in various disorders such as retinopathy, nephropathy, atherosclerosis, neuropathy, *etc*. (31). AGEs accumulate in the pyramidal neurons of brain with ageing (32). Accumulation of AGEs and glycated Tau has been reported in the AD brains paired helical filaments (PHFs) as compared to non-demented brain (13). Tau is found to be glycated at its microtubule-binding region and glycation leads to its decreased affinity for microtubules (33,34) (Fig. 1A). The Tau present in the NFTs is observed to be cross-linked, protease resistant and rendered insoluble as a result of AGEs-modification in AD (35-37). Moreover, Tau glycation is modulated in an isoform-dependent manner and glycation along with phosphorylation increases the aggregation propensity of full-length Tau (38). Glycated Tau has also been reported to induce oxidative stress.(39) AGEs also increase Tau phosphorylation *via* up regulation of aspargine endopeptidases, which inactivates the phosphatase PP2a involved in Tau dephosphorylation (40). Thus, glycation affects the function of Tau not only by cross-linking and inferring protease-resistant but also by inducing other abnormal PTMs-like phosphorylation.

**Figure 1.**
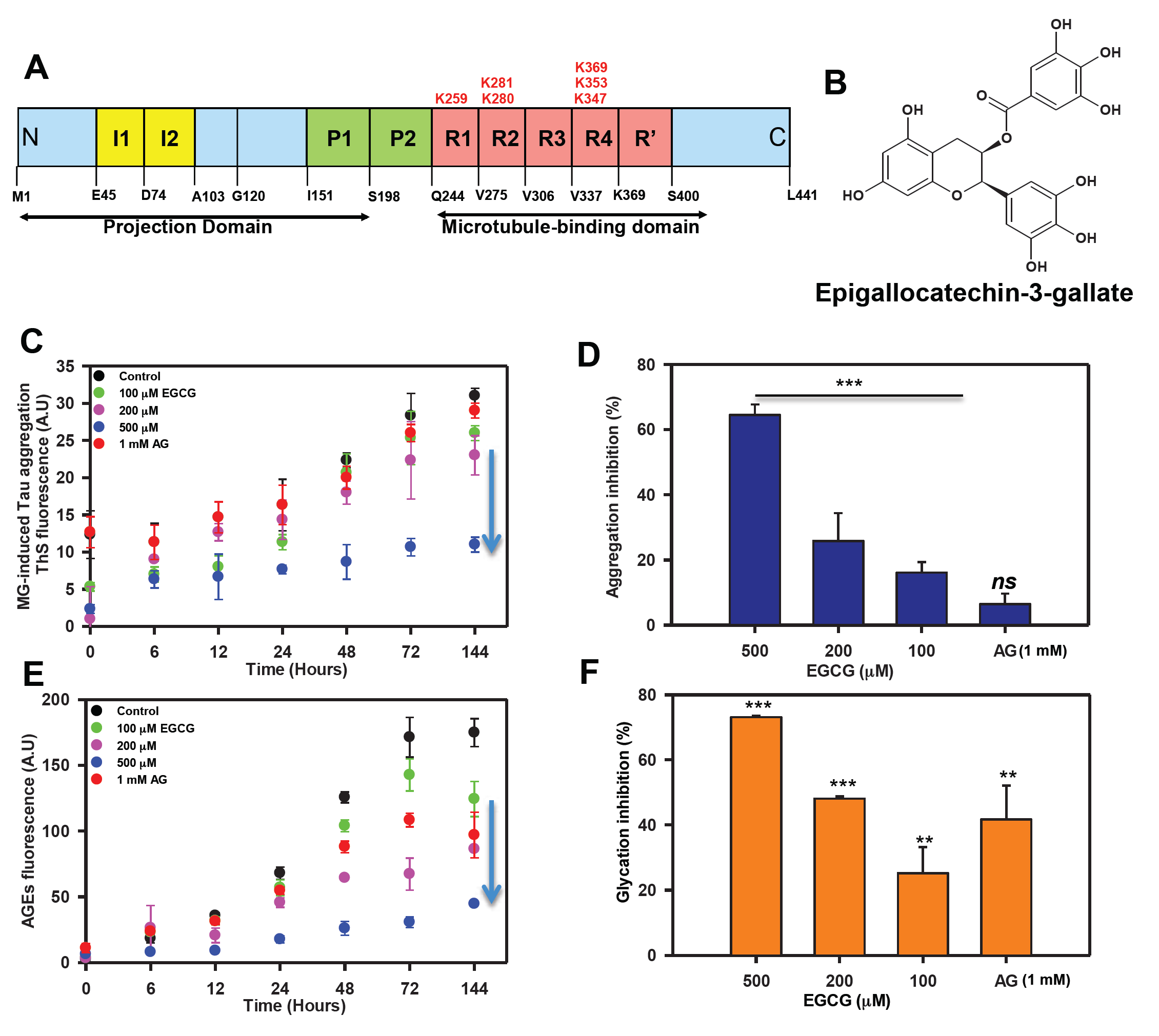
EGCG inhibits Tau glycation. A) Full-length Tau domain organization highlighting the glycation sites. B) Structure of EGCG. C) The ThS analysis of MG-induced Tau aggregation revealed that in absence of EGCG there is increase in ThS fluorescence suggesting aggregation. Increase in concentration of EGCG showed decrease in ThS intensity with lowest at 500 µM of EGCG. The positive control for glycation inhibition, aminoguanidine (AG) showed constant ThS fluorescence. D). The percent inhibition shows EGCG inhibits MG-induced Tau glycation but AG (1mM) does not rescue this aggregation. E) The AGEs auto fluorescence revealed that in presence of EGCG there is less AGEs formation as compared to untreated control. F) EGCG inhibits glycation more efficiently than the positive control AG (1mM) (The values are mean ± std. deviation of two independent experiments. The statistical analysis was carried out by Student’s unpaired T-test with respect to untreated control. *** p≤0.001, **p≤0.01, *p≤0.05. *ns:* non-significant p value).

Cytoskeletal abnormalities of actin and tubulin organization are also being reported in AD. Tau binds and stabilizes microtubules but in modified and diseased state, it looses its affinity for microtubules leading to their destabilization (41,42). Microtubules and the motor proteins play a critical role in vesicular trafficking in neurons, which is an essential neuronal function (43). Along with this, microtubules maintain the neuronal shape as well as they are involved in formation of neuronal growth cones (44). Actin remodeling maintains synaptic plasticity and helps in learning (45).The inclusions of cofilin-actin rods are pathological hallmarks of multiple neurodegenerative diseases. This abnormal actin reorganization is observed in presence of mutant Tau and co-exist with phosphorylated Tau aggregates (46). Moreover, abnormal glycation of actin and tubulin have been reported in the brains of experimental model of diabetes which might play a pathological role in neuronal functioning (47,48).

Epigallocatechin-3-gallate (EGCG) is a green tea polyphenol belonging to sub-class flavonoids (49) (Fig. 1B). EGCG is known to have beneficial effects against cardiovascular diseases (50) and cancer (51,52) and plays a protective role in protein misfolding. EGCG functions in neuroprotection by acting as an anti-oxidant and iron chelator (53). Role of EGCG in modulating Tau post-translational modifications especially glycation is unexplored. In this study, we demonstrate the effect of EGCG in inhibiting Tau glycation *in vitro* and global glycation in the neuroblastoma cells. We report the role of EGCG in rescuing the actin and tubulin cytoskeleton by inhibiting their glycation and thus maintaining neuronal integrity.

## Results

### EGCG inhibits AGE modification of Tau in vitro

Since Tau PTMs like phosphorylation and glycation leads to its altered pathogenic functions, we studied the effect of EGCG on Tau glycation involved in AD pathogenesis. Glycation is non-enzymatic post-translational modification of lysine residues in proteins by sugar molecules and their reactive intermediates leading to protein cross-linking and affecting their structural as well as functional roles. Glycation contributes to AD pathology as it alters and cross-links the protein leading to its aggregation. Moreover, glycation induces formation of reactive oxygen species and EGCG being a known antioxidant, we studied whether it can play a role in glycation. Methyl glyoxal, a reactive intermediate of glucose metabolism was used as a glycating agent. Aminoguanidine (AG) was used as a positive control as it is known to inhibit glycation. We monitored MG-induced Tau aggregation by ThS fluorescence and glycation by advanced glycation endproducts (AGEs) fluorescence. It was observed that EGCG untreated control showed more aggregation as compared to EGCG treated groups (Fig. 1C). 500 µM of EGCG showed 60% inhibition of MG-induced Tau aggregation (Fig. 1D). The AG treated sample did not show decrease in MG-induced aggregation (Fig. 1C, D). The analysis of AGE auto fluorescence revealed that EGCG inhibit glycation of Tau in a concentration dependent manner (Fig. 1E). EGCG was found to be more potent at micro molar concentrations as compared to AG (1 mM). The highest concentration of EGCG 500 µM inhibited AGEs formation by 72% as opposed to 42% by 1 mM AG (positive control) (Fig. 1F).

### EGCG inhibits formation of SDS-resistant Tau AGEs

As EGCG showed a significant inhibition of AGEs in presence of MG in the fluorescence assay, we analyzed these AGEs on SDS-PAGE. The SDS-PAGE analysis showed a decrease in glycated Tau in highest concentration of EGCG (500 µM) at 144 hours of incubation (Fig. 2A, B) as compared to AG. We confirmed the SDS-PAGE results by probing the glycated Tau by AGEs-specific antibody and observed that EGCG treatment inhibits Tau glycation (Fig. 2C). Thus, EGCG acts as potent Tau glycation inhibitor. In order to understand the morphological modulation of Tau AGEs by EGCG, AGEs were visualized by transmission electron microscopy (TEM). The morphological analysis of this glycation revealed the formation of amorphous aggregates by glycated Tau. The AGEs morphology was not altered in presence of EGCG (Fig. 2D). Thus, EGCG inhibits Tau AGEs formation without changes in the morphology of AGEs.

**Figure 2.**
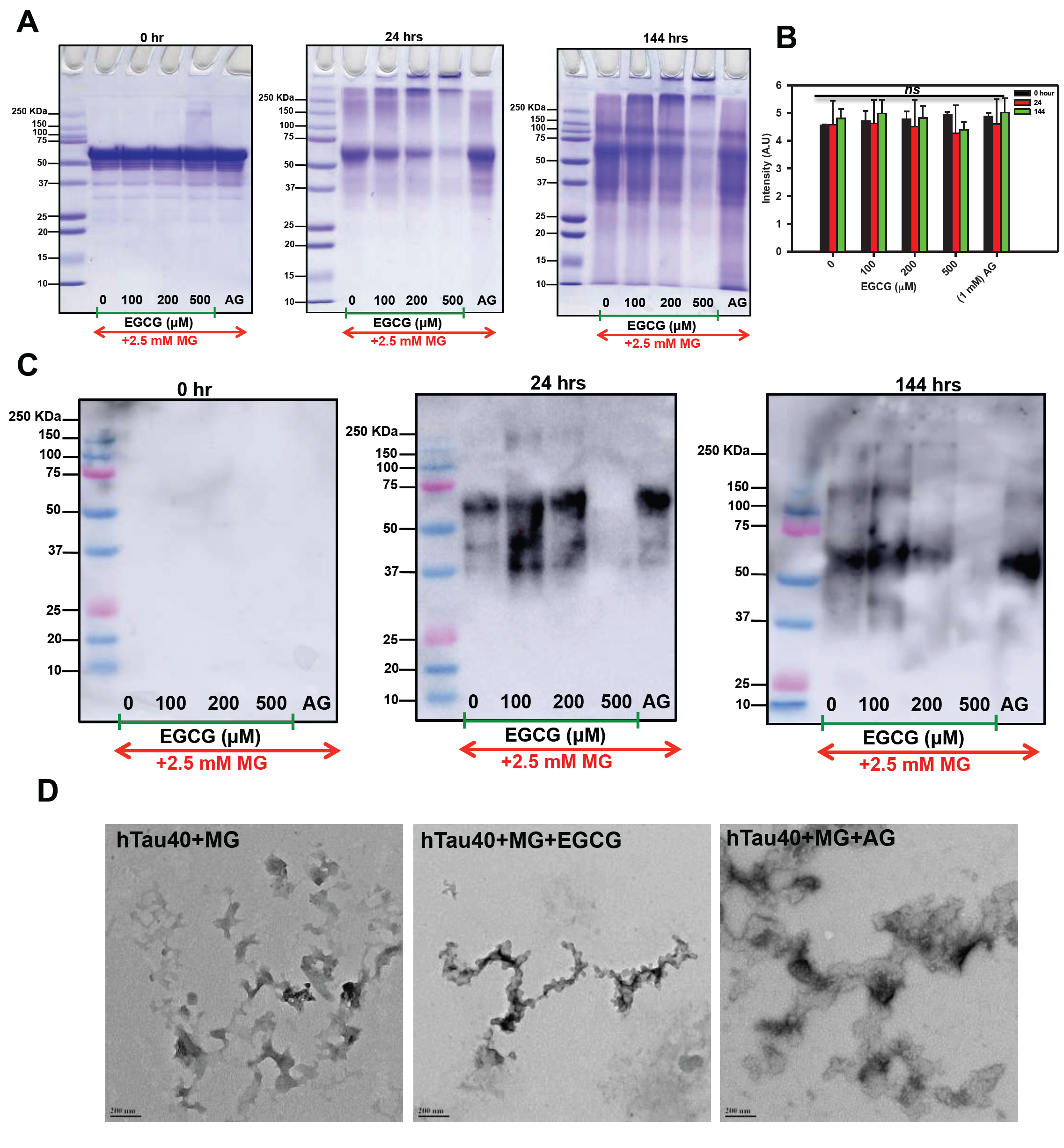
Inhibition of SDS-resistant Tau glycation. A) The SDS-PAGE analysis showed the formation of glycated Tau as smear above the soluble protein band. In presence of EGCG the higher order glycated Tau decreased with increasing concentration and time. B) Quantification of SDS-PAGE shows decrease in intensity in 500 µM EGCG in time dependent manner. C) Immunoblots with AGEs-specific antibody confirm EGCG inhibits Tau glycation in concentration and time dependent manner more efficiently than known glycation inhibitor aminoguanidine. D) The TEM images show presence of amorphous aggregates of glycated Tau and the morphology is not altered by EGCG treatment.

### EGCG inhibits MG-induced AGEs formation and modulates Tau phosphorylation in neuroblastoma cells

EGCG was found potent *in vitro* in inhibiting glycation; we studied its effect on MG-induced AGEs formation in neuroblastoma cells (Fig. 3A). Untreated and EGCG treated cells showed basal level of AGEs as compared to MG treated cells, which showed increased AGEs formation. This increased AGEs formation by MG was reduced in presence of EGCG (Fig. 3B, C). To confirm this observation, immunofluorescence studies were performed for MG-induced AGEs and Tau phosphorylation. AGEs fluorescence was uniform and in the cytoplasm of the cells (Fig. S1A, S2). The cells treated with EGCG along with MG showed basal level of AGEs formation suggesting inhibition of glycation by MG (Fig. 3D). The quantification of fluorescence intensity revealed that MG treatment increased the overall AGEs formation in neuro2a cells, which was attenuated in presence of MG (Fig. 3F). Thus, EGCG shows potent anti-glycation effect. Treatment of MG is also known to induce Tau phosphorylation (54). We studied the effect of EGCG on MG-induced Tau phosphorylation (Fig. 2A). The untreated cells showed basal level of AT100 phosho-Tau distributed evenly throughout cells. EGCG treatment showed distinct nuclear localization of AT100 Tau surrounding the periphery (Fig. 3E zoom, S1B). MG treatment showed nuclear localization but not specifically to the periphery. On treating the cells with MG in presence of EGCG, the nuclear peripheral localization of AT100 was observed in a population of cells (Fig. 3E). The orthogonal cross-sectional analysis confirmed the presence of AT100 in the nucleus surrounding the periphery (Fig. S3). The fluorescence intensity quantification revealed that EGCG enhanced the AT100 Tau in the nucleus as compared to control and MG treated cells (Fig. 3G).

**Figure 3.**
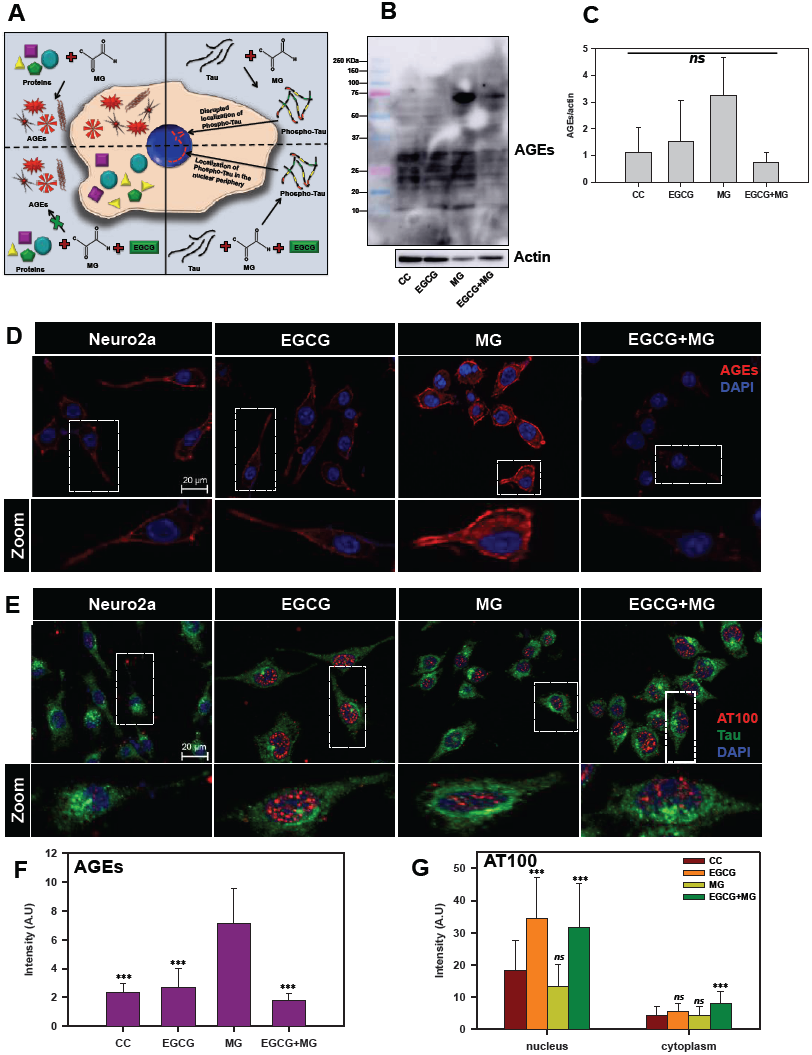
EGCG inhibits MG-induced AGEs formation in neuronal cells. A) A hypothetical model proposing effect of EGCG on MG-induced AGEs formation and Tau phosphorylation. Left panel depicts the effect of MG in inducing AGEs formation, which is abolished in presence of EGCG; the right panel shows the MG-induced Tau phosphorylation and its modulation by EGCG. B) Western blotting of neuronal cells treated with MG for AGEs shows increased AGEs formation in presence of MG as compared to untreated and EGCG treated. The level of glycation by MG is observed to decrease in presence of EGCG. C) Quantification of AGEs formation normalized to actin levels show increased AGEs in MG treated cells as compared to other treatment groups. D) Untreated and EGCG treated cells show basal level of AGEs. MG-induced cells show increased intensity for AGEs. EGCG show decrease in AGEs induced by MG. E) AT100 Tau phospho-epitope is present in negligible levels in control cells and distributed uniformly. EGCG treatment shows a distinct nuclear localization of AT100 Tau around the periphery, which is disrupted in presence of MG. EGCG along with MG shows to restore this AT100 distribution in the nucleus. F) Quantification for AGEs fluorescence confirms increased glycation in MG treated cells and rescue in MG treated cells supplemented with EGCG. G) Tau phosphorylated at AT100 epitope shows enhanced nuclear localization in EGCG treated cells as compared to other groups. (The values are mean ± std. deviation of two independent experiments. The statistical analysis was carried out by Student’s unpaired T- test with respect to MG treatment group. *** p≤0.001, **p≤0.01, *p≤0.05. *ns:* non-significant p value).

### EGCG enhances neuritic extensions

Actin rich neuritic extensions are the key morphological features of neuronal cells. Neuritic extensions are involved in neuronal communication and differentiation. During the initial experiments we observed MG treatment affected neuronal cell morphology leading to rounding off of the cells and loss of neuronal extensions as compared to untreated and EGCG treated cells. Hence, we checked the effect of EGCG on actin cytoskeleton and thus neuritic extensions. We captured Z-stacks and obtained a cumulative 3D image for each group. We observed that EGCG treated cells enhanced neuritic extensions as compared to control cells (Fig. 4A). MG treatment inhibited the neuritic extensions and disrupted actin cytoskeleton in some of the observed cells (Fig S4). Treatment of cells together with MG and EGCG lead to rescue of actin cytoskeleton restoring neuritic extensions (Fig. 4B) As EGCG showed potency in rescuing and enhancing neuritic extensions by remodeling actin cytoskeleton, we checked its effect on microtubule stability since they are critical in maintaining neuronal shape and transport. As with actin, MG also disrupted the microtubule network (Fig. 5A). But EGCG maintained the stability of microtubule network and maintained the neuronal morphology disrupted by MG (Fig. 5A).

**Figure 4.**
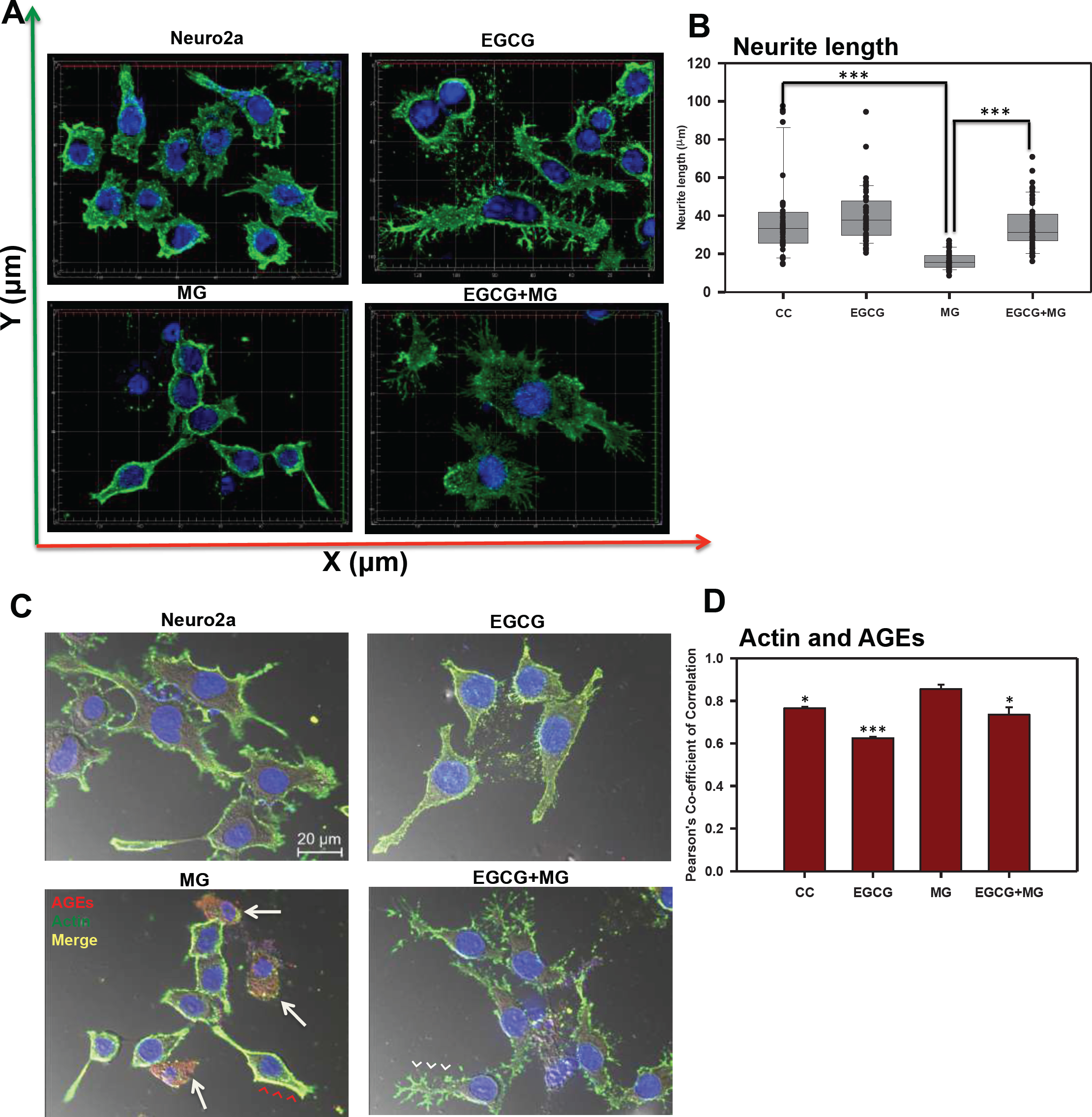
Effect of EGCG on actin cytoskeleton remodeling. A) 3D analysis of actin cytoskeleton shows neuritic extensions, which are maintained in control cells and enhanced in EGCG treated cells. MG disrupts the neuritic extensions severely whereas EGCG restores the actin-rich neuritic extenstions. B) MG treatment drastically reduced the neuritic extensions in cells as compared to control. EGCG treatment rescued the MG affected neurites and maintained the neuronal cell morphology. C) The actin cytoskeleton is severly glycated in presence of MG which is visualized as yellow fluorescence (red arrowheads). The cells with completely disrupted actin cytoskeleton show high levels of AGEs cytoskeleton (white arrows). Alternatively, EGCG together with MG showed decreased actin glycation (white arrowheads) and restoration of the extensions. D) The Pearson’s co-efficient of correlation for co-localization analysis to study the actin glycation by MG showed a higher value as compared to untreated control and EGCG. EGCG was found to decrease the PCC in presence of MG suggesting less colocalization and inhibition of MG-induced glycation. The statistical analysis was carried out by Student’s unpaired T-test with respect to MG treatment group. *** p≤0.001, **p≤0.01, *p≤0.05. *ns:* non-significant p value).

**Figure 5.**
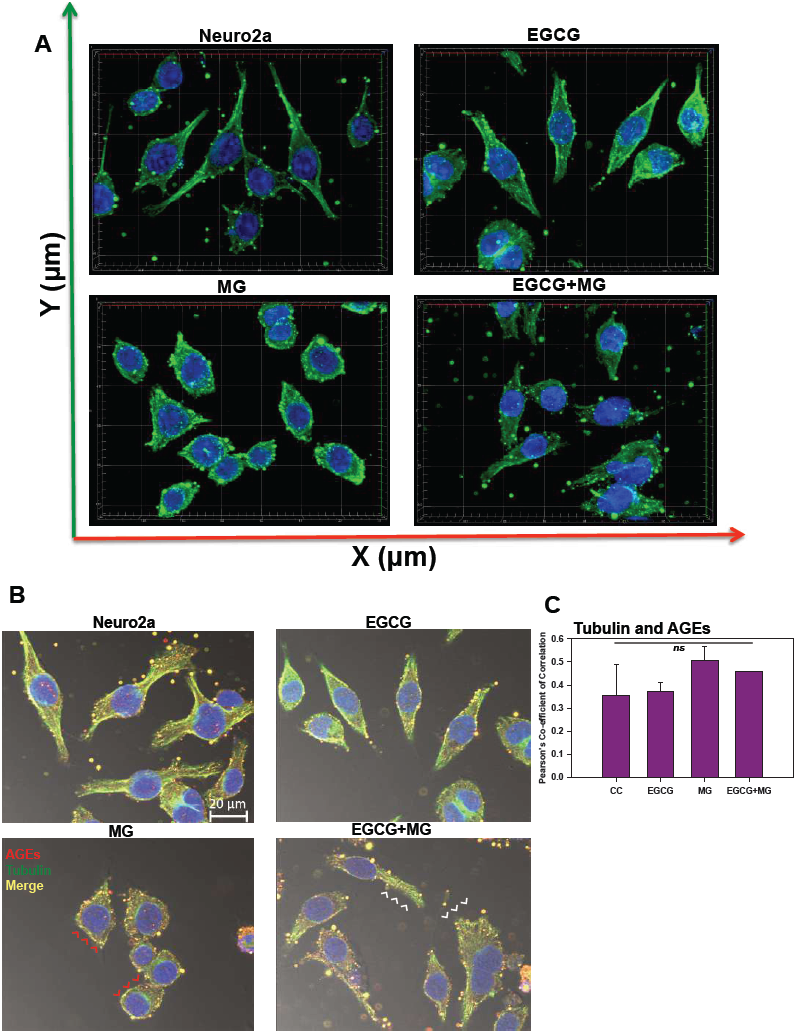
Effect of EGCG on tubulin cytoskeleton. A) Tubulin cytoskeleton was maintained in control and EGCG treated cells with intact MTOC. MG treated cells did not show severe disruption of microtubules but it lead to loss of MTOC integrity. EGCG re-established the integrity of microtubules and MTOC. B) The control and EGCG treated cells showed intact tubulin cytoskeleton with basal levels of AGEs. MG-induced the glycation of tubulin cytoskeleton and lead to its disruption (red arrowheads). On complementation with EGCG, the glycation was inhibited (white arrowheads) and the intact morphology of cells was preserved. C) The PCC for tubulin and AGEs co-localization did not show significant changes among the groups. (The values are mean ± std. deviation. The statistical analysis was carried out by Student’s unpaired T-test with respect to MG treatment group. *** p≤0.001, **p≤0.01, *p≤0.05. *ns:* non-significant p value).

### EGCG inhibits glycation of actin and tubulin cytoskeleton

Since EGCG restored neuritic extensions disrupted by MG, we further studied its effect on preventing glycation of actin and tubulin cytoskeleton. The actin, tubulin and AGEs were mapped by anti-Beta-actin, anti-alpha Tubulin antibody (1:400) and Anti-AGEs antibody (1:200) dilutions respectively. MG-induced glycation of actin and tubulin was observed as yellow fluorescence (Fig. 4C, 5B red arrows heads). Glycation of actin/tubulin cytoskeleton was prevented in presence of EGCG (Fig. 4C, 5B white arrow heads). Moreover, the cells with completely disrupted actin cytoskeleton in MG treated cells showed excessive accumulation of AGEs (Fig. 4B, white arrows). We performed the co-localization analysis for AGEs+actin and AGEs+tubulin for glycation by *coloc2* plugin in Fiji on the background subtracted images (55). It was observed that Pearson’s Co-Correlation Co-efficient (PCC) for actin+AGEs was increased in presence of MG suggesting actin glycation. In presence of EGCG+MG, the PCC decreased confirming the decreased glycation of actin (Fig. 4D). The PCC did not vary significantly for AGEs+tubulin co-localization analysis (Fig. 5C). Thus, EGCG restores cytoskeleton and neuritic extensions by preventing its glycation.

### EGCG maintains cytoskeleton

Actin forms contractile and protrusive ends of the cell maintaining the morphology as well as aiding in cell migration whereas microtubule forms a more polarized network involved in cellular trafficking. B-actin is more abundant in the cell cortex as opposed to tubulin, which is localized to the interior. We demonstrated that this normal distribution of actin and microtubule is disrupted on MG treatment with loss of extensions (Fig. 6A white arrows). The part of cell was focused to make the distribution visible (Fig. 6B, C). EGCG helped maintain this cytoskeleton distribution throughout the cells and inhibited the cytoskeleton disruption by MG. MG differentially affected the two cytoskeletal elements. The actin-rich extensions were severely affected and reduced to small peripheral distribution (Fig. 6). In order to have enhanced view of the cytoskeleton localization we made 3D images from the Z-stacks (Fig. 6D). The untreated and EGCG treated groups showed the peripheral actin and interior tubulin (Fig. 6D zoom). The distribution was hampered in MG treated cells whereas EGCG inhibited the effect of MG and maintained the cell cytoskeleton. In order to get the detailed structural distribution of cytoskeleton elements super-resolution SIM (structured illumination microscopy) was performed. The SIM images re-iterated the results wherein detailed intact distribution of actin and tubulin cytoskeleton was observed in control and EGCG treated neuro2a cells. The focused regions of interest (ROIs) revealed severe disruption of actin extensions and MTOC upon MG treatment. EGCG prevented this disruption and maintained the actin extensions and tubulin network intact. The SIM analyses demonstrated that microtubule network, though not as severely affected as actin, showed loss of microtubule-organizing center in presence of MG (Fig. 7).

**Figure 6.**
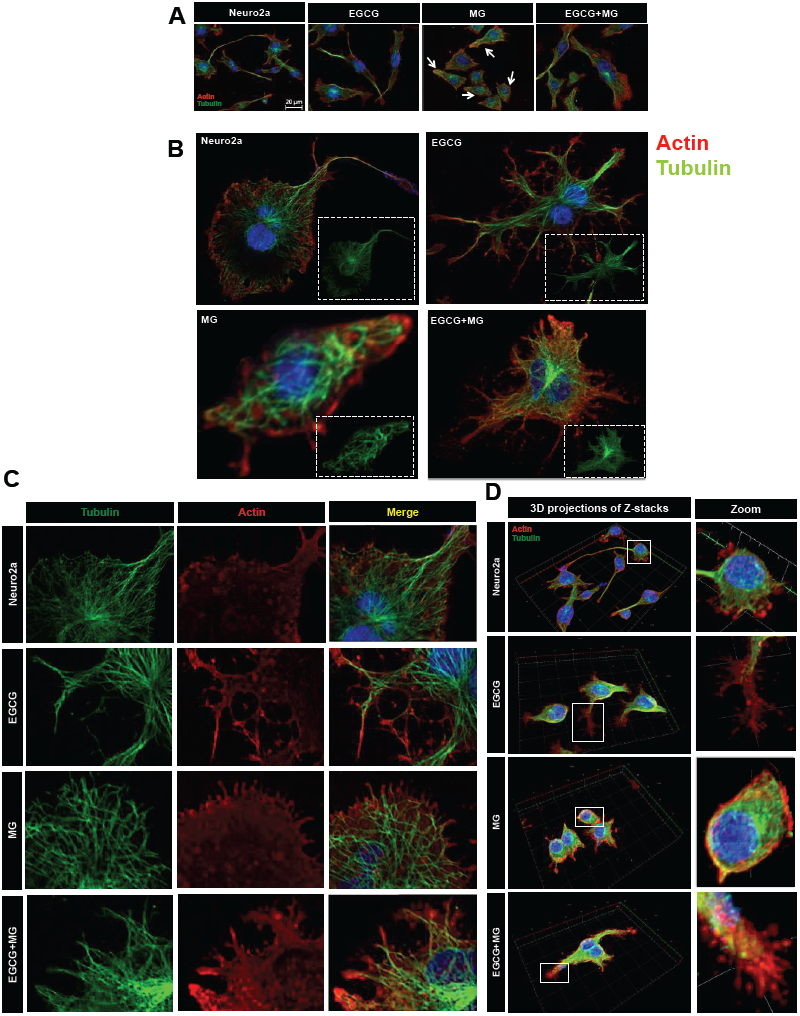
Distribution of cytoskeletal elements. A) Complete cell image showing multiple cells in various treatment groups. Untreated and EGCG treated cells maintain the cytoskeleton structure. MG treated all the cells show loss of cytoskeleton integrity and distribution whereas EGCG treated cells show maintained cytoskeleton distribution and restored integrity. B) Single cell image of the same treatment groups with detailed distribution of cytoskeleton. C) The actin and tubulin cytoskeleton are differentially distributed with actin enriched in small neuritic extensions followed by tubulin fibres. This distribution was found to be hampered on MG treated with excess loss of actin extensions and loss of MTOC integrity. This was unhampered in on EGCG treatment along with MG. D) The 3D analysis of the distribution of cytoskeletal elements shows clear cortical actin as opposed to tubulin in the cytoskeleton. The control and EGCG treated cells shows presence of minute actin extensions protruding out which might be involved in their motility towards other neurons in the vicinity. MG treated cells show loss of these minute extensions. EGCG together with MG showed restoration of these minute actin extensions.

**Figure 7.**
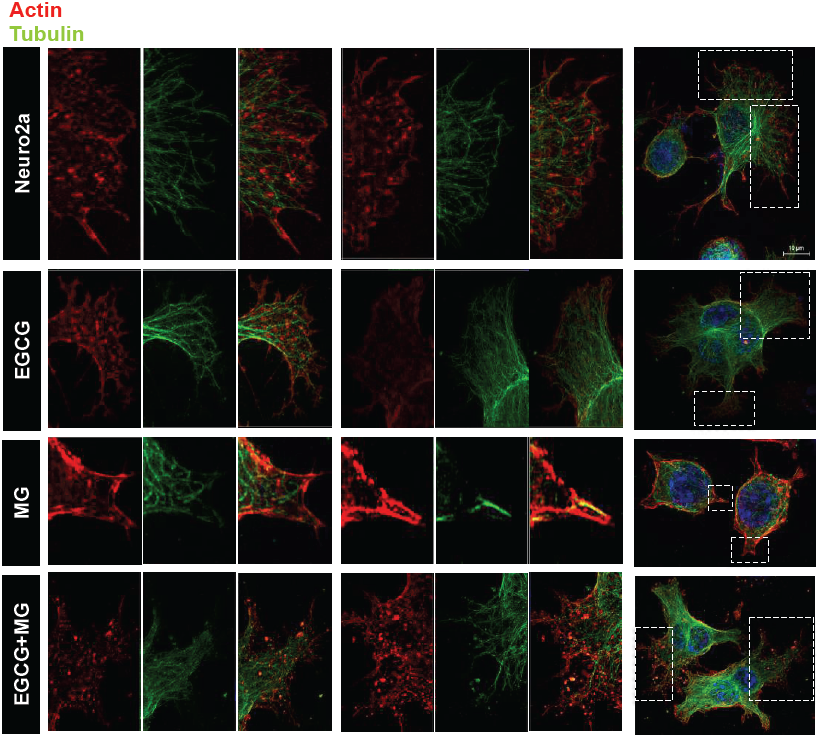
Super-resolution microscopy images for detailed analysis of cytoskeleton. The detailed analysis of distribution of actin and tubulin cytoskeleton was carried out by structured illumination microscopy. The SIM images distinct distribution of actin and tubulin cytoskeleton in control and EGCG treated cells. MG disrupted actin cytoskeleton severely but EGCG prevented this disruption and maintained the cytoskeleton integrity including MTOC.

### EGCG maintains microtubule stability via EB1

Since, microtubule tread milling is one of the essential characteristic, and growth of microtubule ends is aided by +TIPs (microtubule plus end binding proteins) we checked the localization of microtubules with EB1, a +TIP. The overall cells showed disturbed microtubule-EB1 localization in MG treated cells (Fig. 8A). The single cell visualization showed that in control as well as EGCG treated cells; EB1 was localized at the ends of the microtubules (Fig. 8B inset) in the extensions (Fig. 8B). In MG treatment, this localization was found to be disrupted and less abundant. The 3D projections supported this observation (Fig. 8C). But the wide-field fluorescence microscopy imaging was not able to resolve the tubulin-EB1 localization. In order to get an in-depth insight we used super-resolution microscopy. The SIM images gave a clear evidence of loss of tubulin-EB1 localization on MG treatment (Fig. 9). The decrease in abundance of EB1 localization at the ends of the microtubules was also observed. On the contrary, cells treated together with MG and EGCG showed enhanced EB1 at the growing ends of the microtubules. This is observed as the yellow fluorescence for colocalization of tubulin and EB1. Thus, super-resolved images suggest that EGCG may act through EB1 for microtubule stabilization and maintain the structural integrity of the cells. Thus, EGCG may act as enhancer of actin rich extensions and stabilizer of microtubules.

**Figure 8.**
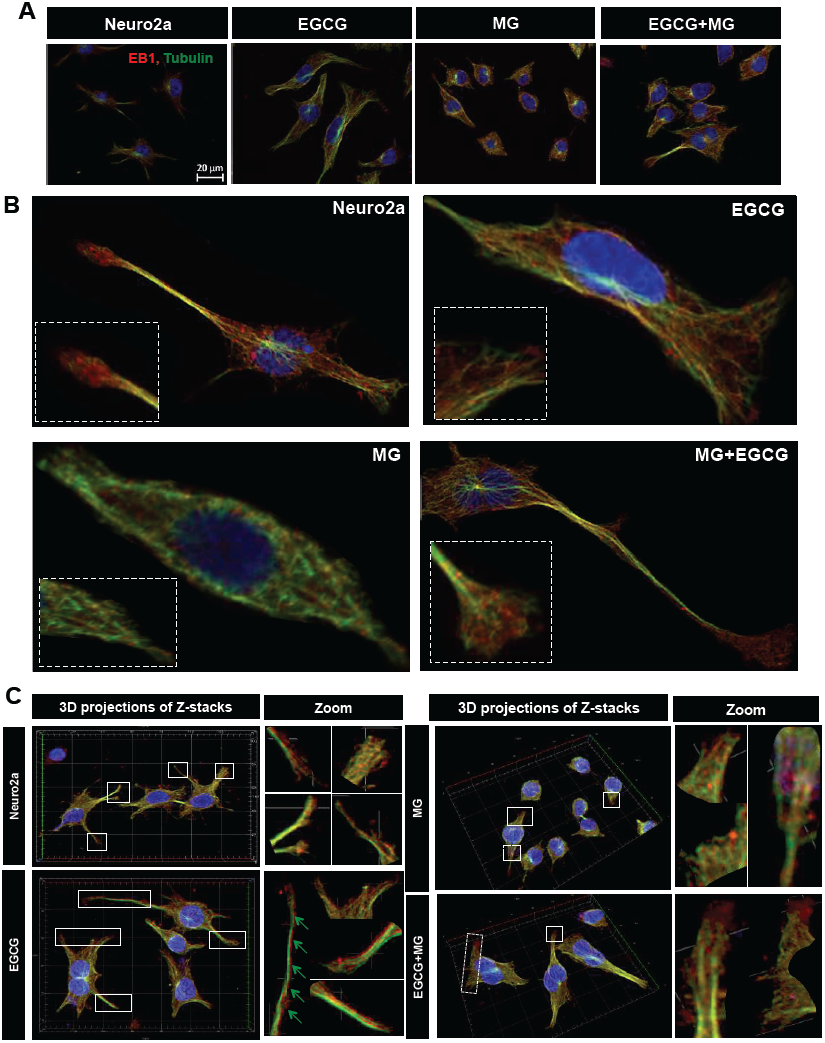
Microtubule stabilization by EB1 maintained by EGCG. A) Cell images showing distribution of EB1 with microtubules in different treatment groups. B) Single cell images showing EB1 maintains the growing end of microtubules helping in its tread milling which is important for cellular functions. EB1 is bound to microtubules (inset) at the ends of the extension but this is not observed in case of MG treatment. MG treatment shows irregular distribution of EB1, which is maintained and resumed by EGCG. The inset enlarged images show the localization of EB1 at the growing ends of microtubules but disrupted by MG. **C)** 3D images of Z-stacks show distinct distribution of EB1 protein at the ends of the microtubules (zoom). MG treatment disrupts the microtubules and hence the localization of EB1. EGCG maintains the microtubule EB1 localization in presence of MG thus stabilizing the microtubules.

**Figure 9.**
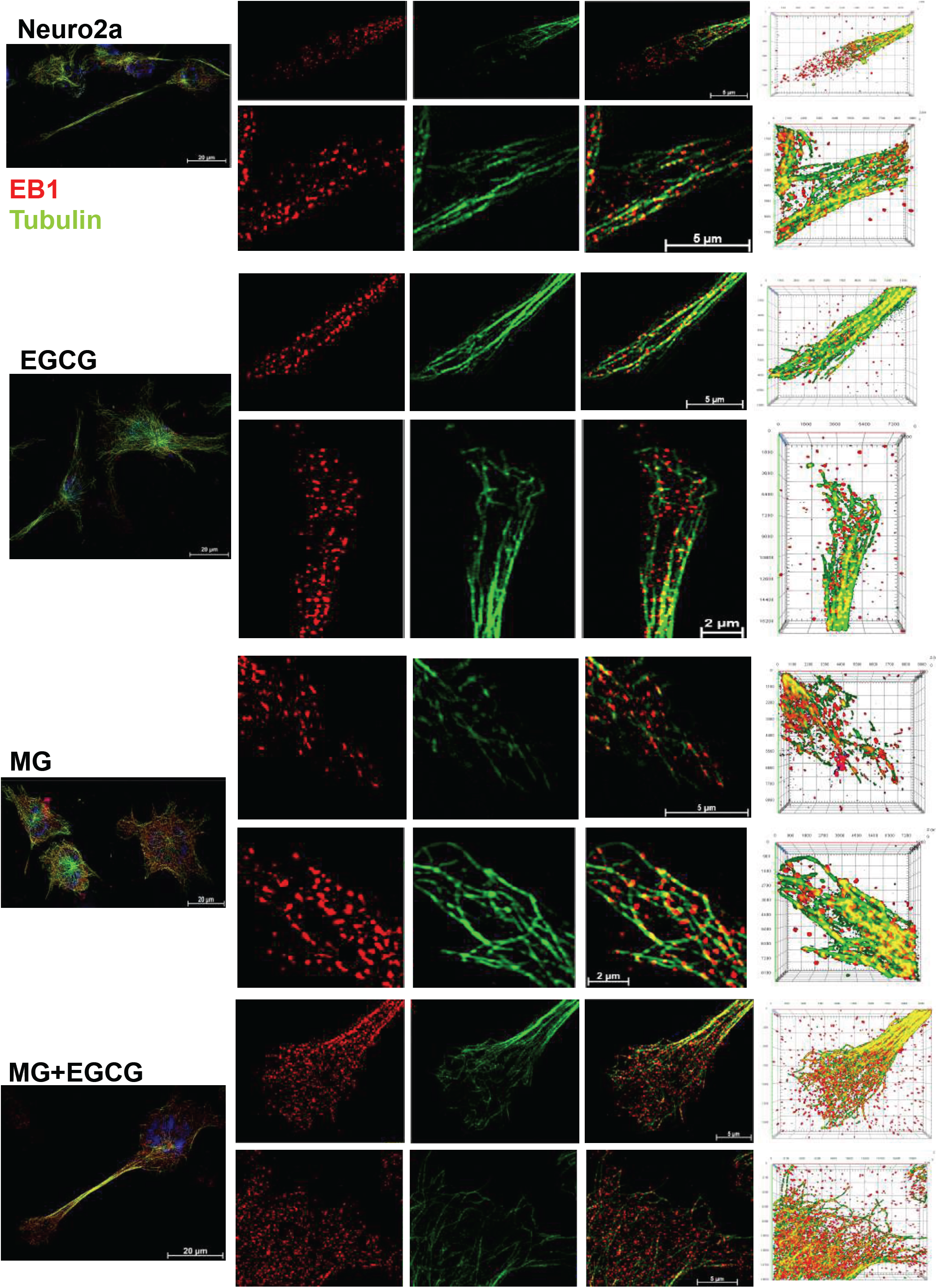
Super-resolved images for EB1-mediated microtubule stabilization. The SIM images reveal clear localization of EB1 at the ends of microtubules in the control and EGCG treated cells. The EB1 localization was severely hampered and less abundant in MG treated cells. This effect of MG is inhibited in presence of EGCG. The rescued colocalization is visualized as yellow fluorescence.

**Figure 10.**
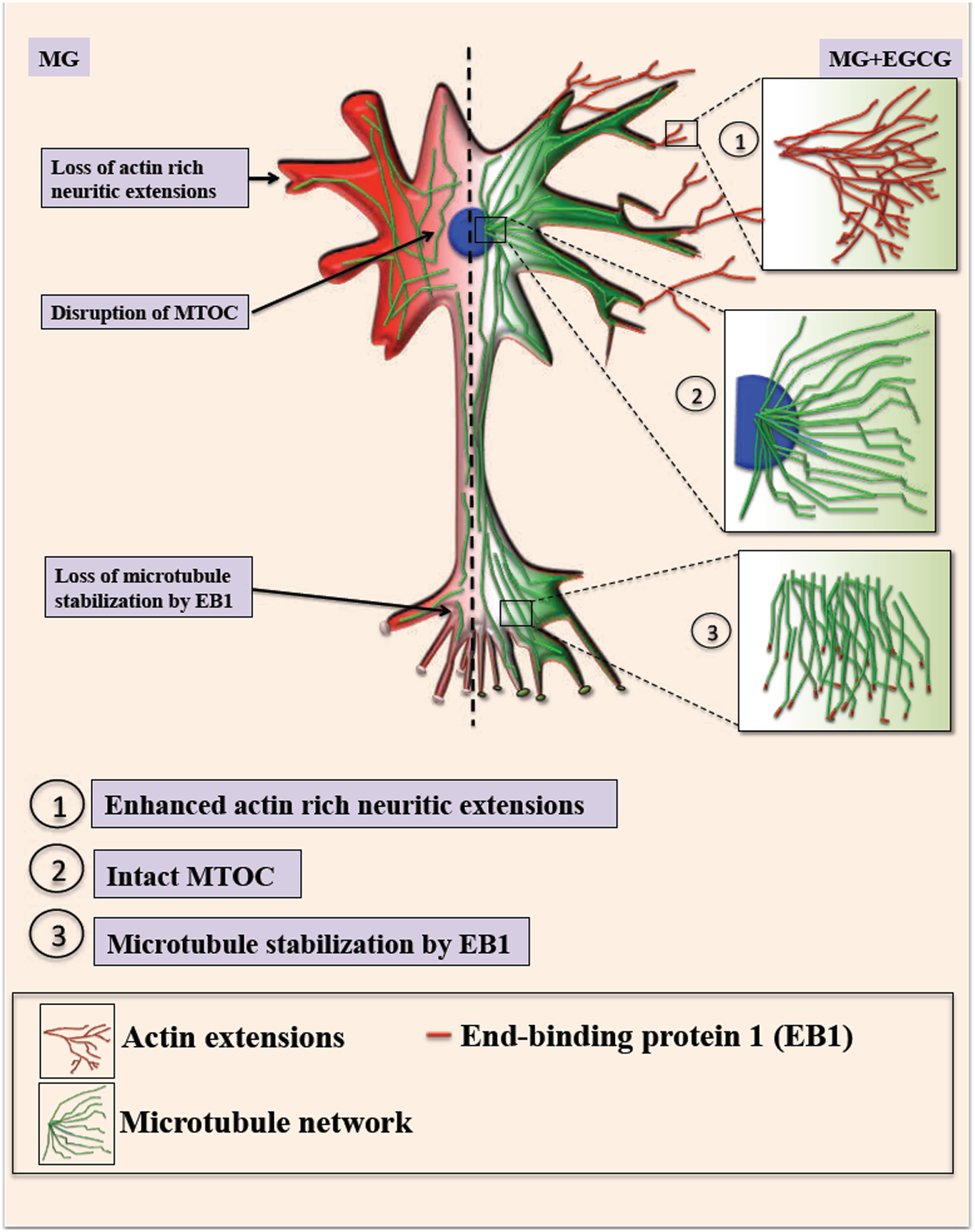
EGCG maintains neuronal cell cytoskeleton integrity. Methyl glyoxal treatment leads to glycation of actin and microtubules leading to hampered growth of neuritic extensions and microtubule organization. Microtubule stabilization by +TIP EB1 is also affected resulting in loss of cell morphology. In presence of EGCG, the actin rich neuritic extensions are enhanced which might help in neuronal connections. EGCG also maintains intact MTOC and thus microtubule mediated transport. EGCG also helps in microtubule polymerization *via* EB1 protein thus maintaining and rescuing overall cell integrity.

## Discussion

The abnormal post-translational modifications of Tau impacts are cellular localization and functioning in the neuronal cells. Though, voluminous studies report pathological effects of Tau hyperphosphorylation and its inhibition by specific molecules like kinase inhibitors (56), enhancers of phosphatase activity (57), Tau acetylation inhibitors (15) *etc*., complete rescue of Tau pathology is yet to be studied. Glycation on the other hand, is not a major Tau modification, but is known to enhance Tau pathology by stable cross-linking, protease resistance and inferring insolubility. Moreover, Tau glycation has already been known to lead to its aggregation and abolish microtubule-binding function. The current study reports a potent anti-glycation activity of EGCG on inhibiting MG-induced Tau glycation. EGCG modified with fatty acids have been shown to posses anti-glycation activity *in vitro* (58). EGCG is also known to posses the carbonyl trapping property with the ratio of 3 moles of EGCG per mole of MG (59). In our studies, EGCG was found to be more potent than the known glycation inhibitor aminoguanidine. EGCG potently inhibited glycation with 5 fold excess of MG whereas AG showed inhibition with 2.5 excess of MG suggesting EGCG is twice as potent than AG in inhibiting Tau glycation.

EGCG potently rescued MG-induced AGEs formation in neuroblastoma cells. The significant decrease in MG-AGEs was observed in neuro2a. Similar studies have reported decrease in MG-induced AGEs formation in neuro2a by AG (54). The same study demonstrates MG to induce Tau hyperphosphorylation and up regulate GSK3β. We have similar observations wherein MG disrupted the unique nuclear localization of Tau AT100, which was observed in EGCG treated cells. The exact significance of this unique localization still needs to be explored. Thus, EGCG can modulate dual modifications of glycation and phosphorylation-induced by MG. The decrease in AGEs and ROS by EGCG has already been shown by previous studies *via* Nrf2 (Nuclear factor erythroid 2-related factor 2)-dependent pathways in various cell types (60). The exact mechanism of inhibiting AGEs formation in mouse neuroblastoma cells is unclear but it might be due to the quenching of carbonyls formed during glycation by EGCG. The cytoskeletal proteins like actin and tubulin play a crucial role in maintaining the axonal integrity and fast axonal transport (FAT) (61,62). Tubulin glycation have been reported in diabetic rat model affecting the axonal transport (63). Moreover, the reactive lysines are essential for microtubule assembly, which are modified by glycation (64,65). Thus, glycation might affect the tubulin assembly. Glycation of actin decreases its polymerization into F-actin (66), which plays a role in stabilizing spine morphology (67). EGCG was found to reduce the glycation of cytoskeletal elements like actin and tubulin. Reduced glycation was accompanied with resumed tubulin assembly and enhanced actin extensions thus improving neuronal structural integrity (Fig. 9). Moreover, tubulin stabilization by EB1 was enhanced in presence of EGCG, which might be *via* interaction of Tau with EB1 (68,69).

## Conclusion

The Tau-targeting therapeutics are being widely studied with respect to their Tau PTMs modifying potential especially phosphorylation. Glycation though not the major modification of Tau but is known to stabilize the preformed Tau PHFs aggravating the pathology. We report, EGCG as a modulator of Tau glycation and inhibitor of AGEs in the neuroblastoma cells. Additionally it helps to maintain the neuronal cytoskeleton by rescuing its modification. Thus, EGCG may be considered as a novel therapeutic targeting Tau PTMs remodeling cytoskeleton integrity.

## Experimental procedures

### Chemicals and reagents

Luria-Bertani broth (Himedia); Ampicillin, NaCl, Phenylmethylsulfonylfluoride (PMSF), MgCl_2,_ APS, DMSO, Ethanol (Mol Bio grade), Chloroform, Isopropanol (Mol Bio grade) were purchased from MP biomedicals; IPTG and Dithiothreitol (DTT) were obtained from Calbiochem; EGCG, MES, BES, SDS, MTT reagent, Okadaic acid and TritonX-100 from Sigma; EGTA, Protease inhibitor cocktail, Tris base, acrylamide and TEMED were purchased from Invitrogen. The cell biology reagents Dulbecco modified eagle’s media (DMEM), Fetal bovine Serum (FBS), Horse serum, Phosphate buffer saline (PBS, cell biology grade), Trypsin-EDTA, Penicillin-streptomycin, Pierce™ LDH Cytotoxicity Assay Kit (Thermo, cat no 88953), RIPA buffer were also purchased from Invitrogen. The uncoated glass coverslips for immunofluorescence studies (18 mm) were purchased from Blue star. The following antibodies were used for the immunofluorescence studies; Anti-AGE (Advanced Glycated End-products) Antibody goat polyclonal 1:200 (Millipore Cat. No. AB9890) Beta-actin loading control 1:400 (Thermo fisher cat no. MA515739) rabbit alpha Tubulin 1:400 (Abcam, cat no ab176560), EB1 Monoclonal Antibody (KT51) Thermo fisher cat no. MA172531) total Tau antibody K9JA 1:500 (Dako, cat no A0024), anti-mouse secondary antibody conjugated with Alexa flour-488 1:500 (Invitrogen, cat no A-11001), Goat anti-Rabbit IgG (H+L) Cross-Adsorbed Secondary Antibody with Alexa Fluor 555 1:1000 (A-21428), Rabbit anti-Goat IgG (H+L) Cross-Adsorbed Secondary Antibody with Alexa Fluor 594 (A27016) 1:500 and DAPI (Invitrogen).

### Tau purification

Protein purification for full-length Tau was carried out as previously described (70). In brief, full-length recombinant Tau was expressed in *E*.*coli* BL21* strain. The cell pellets were homogenized under high pressure (15,000 psi) in a microfluidics device for 15 minutes. 0.5 M NaCl and 5 mM DTT was added to the lysate befor heating at 90°C for 15 minutes. The cooled lysate was centrifuged at 40,000 rpm for 50 minutes. The supernatant dialyzed in Sepharose A buffer overnight. Further clarification of lysate was performed by centrifuging at 40,000 rpm for 50 minutes. This pre-clarified lysate was subjected to cation-exchange chromatography.(Sepharose fast flow GE healthcare) for further purification. The fractions containing Tau were pooled and subjected size exclusion chromatography (16/600 Superdex 75pg GE healthcare). The Tau concentration was measured using BCA method.

### Glycation inhibition assay

20 µM Tau was incubated with 2.5 mM of methyl glyoxal (MG) in BES buffer (pH 7.4), 25 mM NaCl 1mM DTT, protease inhibitor cocktail, 0.01% Sodium azide, and different concentrations of EGCG 100, 200, 500 µM. Ammonium guanidine (1 mM) was used as a positive control for glycation inhibition. The reaction mixtures were incubated at 37°C and monitored regularly for aggregation by ThS fluorescence. The advanced glycation end products were monitored by auto fluorescence excitation 370 nm and emission 432 nm. The readings were obtained in triplicates for 2 sets of experiments.

### Electron microscopy

MG-induced Tau aggregates in presence and absence of EGCG were visualized by Transmission electron microscopy. The reaction mixtures were diluted to 2 µM and applied to 400 mesh carbon coated copper grids for 1 minute. The grids were washed twice with filtered Milli Q water for 45 seconds each. The grids were further incubated with 2% uranyl acetate. The samples were scanned using Tecnai G2 20 S-Twin transmission electron microscope. Experiments were performed for 2 sets of data.

### SDS-PAGE

The samples from reaction mixtures made for glycation in presence and absence of EGCG were separated on 10% SDS-PAGE at 0, 24 and 144 hours and stained with Coomassie brilliant blue stain.

### Immunoblotting

Tau glycation inhibition by EGCG was confirmed by immunoblotting the reaction mixtures at intervals of 0, 24 and 144 hours with AGEs-specific antibody. The reaction mixtures at the given time points were separated on 10% SDS PAGE and transferred onto the methanol-activated PVDF membrane. The membrane was blocked in 5% BSA for 1 hour and probed with anti-AGEs (1:2000 dilutions) antibody overnight at 4°C. The unbound primary antibody was removed by 3 washes of PBST and the blot was incubated with secondary antibody donkey anti-goat HRP (1:5000 dilutions). The blot was developed by using ECL Plus chemiluminiscence kit.

For the cell biology studies, the cells were seeded as 1.5×10^5^ for 24 hours in DMEM supplemented with 10% FBS and antibiotics penicillin-streptomycin. Four experimental groups were maintained as cell control, EGCG (100 µM) treated, MG (1 mM) treated and MG+EGCG treated. Treatment was carried out for 24 hours after which, the cells were lysed in RIPA buffer and 75 µg of lysate was loaded onto the 10% SDS-PAGE. The immunoblotting protocol was carried out as described above for the biochemical studies.

### Immunofluorescence

Neuro2a cells were maintained in Advanced DMEM supplemented with 10% FBS and antibiotics penicillin-streptomycin. For immunofluorescence studies 5×10^4^ cells were seeded on glass coverslip in a 12 well culture plate for 24 hours. Four experimental groups were maintained as cell control, EGCG (100 µM) treated, MG (1 mM) treated and MG+EGCG treated. Treatment was carried out for 24 hours after which, the cells were fixed with ice-cold methanol. Cell permeabilization was carried with 0.2% TritonX 100. After 3 PBS washes cells were blocked in 5% horse serum for 1 hour at 37°C. Cells were incubated with primary antibodies at 4°C overnight. Further, cells were washed with 1X PBS and incubated with respective antibodies anti-mouse secondary antibody conjugated with Alexa flour-488, Goat anti-Rabbit IgG (H+L) Cross-Adsorbed Secondary Antibody with Alexa Fluor 555 (A-21428) followed by 300 nM DAPI. The coverslips were mounted in 80% glycerol and sealed and were observed under 63X oil immersion lens in Axio Observer 7.0 Apotome 2.0 (Zeiss) microscope using ZEN pro software.

### 3D processing of the immunofluorescence images

All the images were acquired using 63X oil immersion lens in Axio Observer 7.0 Apotome 2.0 (Zeiss) microscope. The 3D Z-stack images were obtained with optimum slice size of 0.24 µm. The slice size may vary depending on cell thickness. After acquiring the 3D Z-stacks images were processed for orthogonal projections to demonstrate the cross-sections of the cells. The quantitative analysis of the fluorescence intensities were done for the respective orthogonal projections by ZEN image processing software and the mean intensities were plotted using Sigma Plot 10.2.

### Co-localization analysis

To ensure the glycation of cytoskeleton, co-localization analysis was carried out for AGEs (red) and actin (green) and AGEs (red) tubulin (green) fluorescence. The analysis was carried out on the background subtracted images the reported algorithm (55). The Pearson’s co-efficient of correlation was determined by the *coloc-2 plugin* of Fiji software. The obtained values were plotted in Sigma Plot 10.2 and statistical analyses were carried out.

### Structured Illumination Microscopy

All the super-resolved images were captured on ZEISS Elyra 7 with Lattice SIM microscope. The Z-stacks were obtained and further processed in the Zen imaging software.

### Statistical analysis

All the statistical analyses were carried out by using unpaired T-test by Sigma Plot 10.2. For the *in vitro* Tau glycation experiments the percentage inhibition is calculated with respect to control (EGCG untreated). The error bars represent mean ±SD values. 95% confidence intervals were maintained for the analyses. The quantification of neurite extensions was carried out by neurite tracer plugin of Fiji. An average of 10 images containing multiple cells were analyzed for the neurite length. The determination of Pearson’s co-efficient of correlation for fluorescence colocalization was done by an in-built plugin *coloc-2* of Fiji.

## Abbrevations

PTMs: Post-translational modifications
AGEs: Advanced glycation end-products
MG: Methyl glyoxal
EGCG: Epigallocatechin-3-gallate
MTOC: Microtubule Organizing Center
EB1: Microtubule-associated protein RP/EB family member 1
Nrf2: Nuclear factor erythroid 2-related factor 2
PCC: Pearson’s correlation coefficient.

## Acknowledgements

### Funding

This project is supported in part by grants from the Department of Biotechnology from Neuroscience Task Force (Medical Biotechnology-Human Development & Disease Biology (DBT-HDDB))-BT/PR/19562/MED/122/13/2016 and in-house CSIR-National Chemical Laboratory grant MLP029526. SS acknowledges DBT for the fellowship. The author acknowledges Prof. Mahesh Kulkarni for his valuable comments and fruitful discussion of this manuscript.

### Author contributions

SKS and SC conducted most of the experiments, analyzed the results, and wrote the paper. SC Conceptualization provided resources, formal analysis, supervision, validation and wrote the paper.

---

**Supplementary figure 1.**
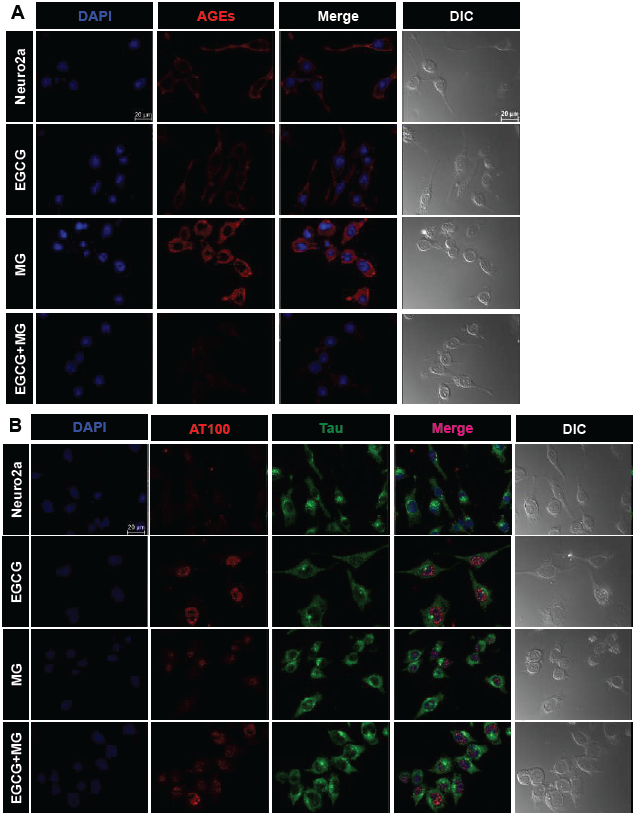
MG-induced glycation and Tau phosphorylation. **A)** Single channel images showing increase in AGEs on MG treatment in neuro2a cells and their inhibition by EGCG. Neuro2a cells untreated and EGCG treated show neuritic extensions and intact cell shape for both modifications. MG treatment alters the cell morphology resulting in loss neuritic extensions and rounding off of cells. Supplementation of EGCG with MG shows normal cell morphology with maintained cell shape and neuritic extensions. B) AT 100 localization on EGCG treatment surrounding nuclear periphery which is disrupted by MG treatment.

**Supplementary figure 2.**
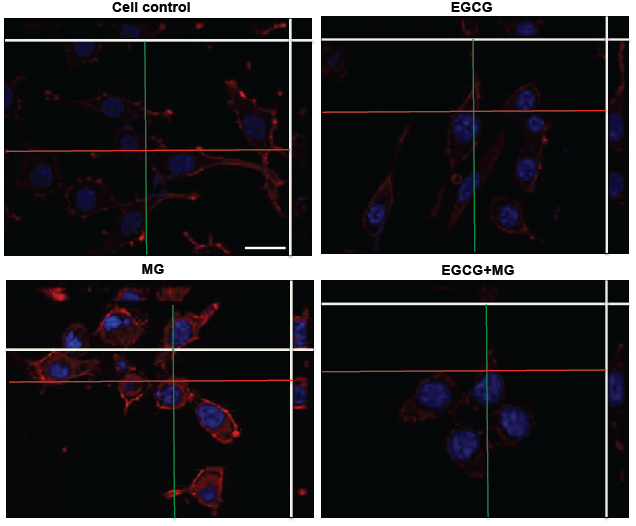
Orthogonal projection analysis of MG-induced AGEs positive neuro2a cells. Orthogonal projectional analysis shows basal level of AGEs in untreated and EGCG treated cells. MG treatment induces global glycation in the cells resulting in enhanced AGEs formation in the cells. EGCG is found to inhibit this effect of MG and reduce the global AGEs formation in the cells. Scale bar: 20 µm.

**Supplementary figure 3.**
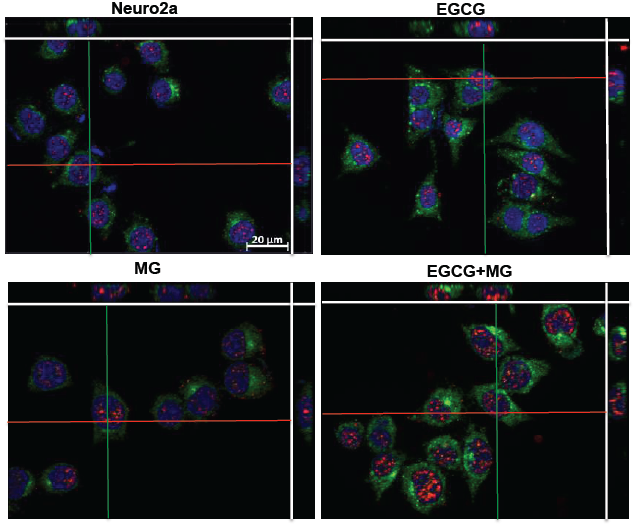
Orthogonal projection analysis of MG-induced Tau phosphorylation in neuro2a cells. AT100 phospho-Tau is present at basal levels in control cells and distributed throughout the cytoplasm and nucleus. EGCG treatment is found to change the localization of phospho-Tau in the nucleus at the periphery in a ring like manner. MG treatment disrupts this arrangement in as seen in the orthogonal projections whereas MG and EGCG together maintain the AT100 phospho-Tau in the nucleus.

**Supplementary figure 4.**
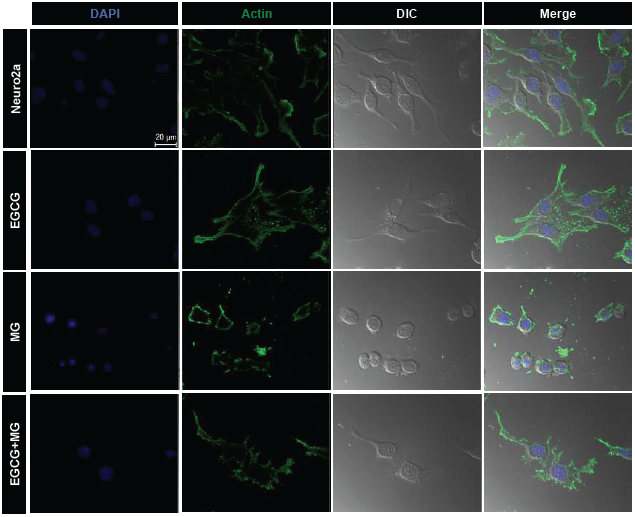
MG-induced disruption of actin cytoskeleton. Untreated neuro2a cells show visible neuritic extensions rich in actin also evidenced by merge of DIC with actin. EGCG treated cells more of minute neuritic extensions along with the long extensions. MG treatment severely disrupts actin cytoskeleton and neuritic extensions (merge). EGCG treatment with MG leads to formation of both minute and long actin rich neuritic extensions.

